# Temperature and host plant ecotype drive nitrogen fixation, but not nodule community composition, in hairy vetch

**DOI:** 10.1101/2024.08.28.610160

**Authors:** Rebecca Fudge, Paula Gardner, R Ford Denison, Liana Burghardt, Julie Grossman

**Author notes:** Corresponding author: Julie Grossman.

## Abstract

Hairy vetch (*Vicia villosa* Roth) is a commonly grown cover crop throughout the U.S., which can contribute nitrogen for subsequent cash crops through biological nitrogen fixation (BNF) in association with *Rhizobium leguminosarum* biovar *viciae* (Rlv) bacteria. Hairy vetch is one of the few cover crops sufficiently cold-tolerant to over-winter in the Upper Midwestern U.S. However, nitrogen contributions by hairy vetch vary across locations, potentially due to cold impacts on the legume/rhizobia symbiosis. The traditional route to improve BNF in legumes involves selecting superior rhizobia strains to create more effective inoculants to apply at planting, but inoculants often fail to compete and survive in agricultural soils. Instead, this study tested the effects of temperature and host plant ecotype on hairy vetch BNF and Rlv community composition in nodules, with the goal of potentially identifying vetch ecotypes able to select beneficial Rlv strains from the soil community. Four hairy vetch ecotypes trapped Rlv from three Minnesota soils, at warm or cold temperatures. Vetch ecotype was a key driver of BNF and nodule formation under warm and cold conditions. However, temperature and plant ecotype did not drive Rlv community composition in nodules, and Rlv community composition did not affect plant productivity. Taken together, these results suggest that the best strategy to improve BNF at low temperatures in hairy vetch likely depends on breeding for improved biomass accumulation and nitrogen fixation in host plants, rather than focusing on host plant selection of beneficial rhizobia.

## 1. Introduction

The legume/rhizobia symbiosis is among the earliest examples of human use of beneficial microbes to enhance crop yields. This mutualism depends on rhizobia soil bacteria, which form organs called nodules on legume roots and fix atmospheric nitrogen into a plant-available form in exchange for carbon resources from the plant (Reviewed in Lindström and Mousavi, 2020). Prior to widespread use of synthetic N fertilizers in the 20^th^ century, this symbiotic exchange, called biological nitrogen fixation (BNF), accounted for almost all new nitrogen entering agroecosystems (Vitousek et al., 2013). While synthetic fertilizers led to breakthroughs in agricultural yields, they also have negative environmental impacts, especially their contribution to water pollution and nitrous oxide emissions (Tilman et al., 2002; Walling and Vaneeckhaute, 2020).

One way to mitigate these negative effects is to increase BNF in a manner that is compatible with U.S. agricultural systems, which are dominated by summer annual field crops (USDA NASS, 2022). Winter annual cover crops are a promising option because they are planted after cash crop harvest in the fall, grown through winter, and terminated and incorporated into soil in spring. Leguminous cover crops contribute biologically fixed nitrogen for use by following cash crops during typically fallow winter months. Hairy vetch (*Vicia villosa* Roth) is one of the most widely grown cover crops in the United States (CTIC et al., 2023) for its winter hardiness, high biomass production, and nitrogen contributions to subsequent cash crops (Brandsæter et al., 2008; Mirsky et al., 2017; Perkus et al., 2022; Power and Zachariassen, 1993). In warmer climates such as North Carolina, hairy vetch can contribute up to 122 – 217 kg N ha^-1^ with over 90% N derived from the atmosphere, compared to 68 - 156 kg N ha^-1^ for crimson clover (*Trifolium incarnatum*), and 67 - 126 kg N ha^-1^ and for Austrian winter pea (*Pisum sativum*) (Parr et al., 2011).

Despite the documented utility of hairy vetch, few improved cultivars exist, and most commercially available varieties experience winter kill in the upper Midwestern U.S., with inconsistent biomass accumulation and BNF (Kucek et al., 2019). New cold-tolerant accessions of hairy vetch have been bred for use in these regions and have improved freezing tolerance (Maul et al., 2011; Wiering et al., 2018), but their nitrogen contributions to following crops remain variable (Perrone et al., 2020). Cold temperatures are known to negatively impact the legume-rhizobia symbiosis by inhibiting legume nodulation and nitrogenase activity Thus, this study focuses on an alternative path that considers the plant-associated microbiome during breeding, with a goal of recruiting beneficial microorganisms (Busby et al., 2017; Nerva et al., 2022; Trivedi et al., 2022). For legumes, this would involve identifying cultivars that may select superior rhizobia strains from the soil environment (Denison, 2021). These superior strains selected by plant hosts could improve BNF efficiency (N2 fixed per CO2 respired or per g nodule mass) to prevent plants from investing too much carbon in nodules at the expense of biomass accumulation.

In fields where legumes have previously been grown, a population of naturalized rhizobia is often established in the soil. These naturalized rhizobia vary in many traits, including their competitiveness for nodulation and nitrogen fixation efficiency (Ballard and Charman, 2000; Drew and Ballard, 2010; Nandasena et al., 2007). When subsequent legumes are grown in soil with a naturalized rhizobia population, they selectively associate with some, but not all, rhizobia strains present. As a result, plant host genotype has been found to be a driver of nodule microbiome assembly (Brown et al., 2020; Burghardt et al., 2022, 2018). In hairy vetch inoculated with soil from three locations, plant genotype accounted for 12.5% of diversity among rhizobia strains, even as soil origin—especially variation in pH and phosphorus—explained the most variation in rhizobia diversity (N.V. Mothapo et al., 2013). Other studies have investigated whether the nodule microbiome selected by certain host plant genotypes offers more plant growth benefits. Genotypes of the reference legume *Medicago truncatula* differed in their selection of rhizobia communities, with genotype A17 preferentially supporting nodules containing strains that, in single-strain inoculation, led to faster biomass growth (Burghardt et al., 2018). In addition to current benefits, host plants that selectively support more-beneficial strains could enrich the soil with those strains.

Another selective force acting on soil rhizobia populations is temperature. In a study of rhizobia evolution in the absence of plant hosts (Burghardt et al., unpublished data), soil rhizobia populations subjected to cold temperatures experienced reduced strain diversity compared to those in soil held at ambient temperatures. Specifically, cold conditions altered the strain community to be more beneficial for one *Medicago truncatula* genotype, resulting in increased vegetative biomass, yet less beneficial for a different genotype. Separately, a study of soybean showed that while the rhizobia species *B. japonicum* and *B. elkanii* formed equal numbers of nodules on soybean at ambient temperatures, *B. japonicum* occupied the vast majority of nodules at low temperatures (Suzuki et al., 2014). Taken together, these studies suggest that low temperatures affect rhizobia fitness in soil, and/or strains’ competitiveness for nodulation.

The most common time for hairy vetch planting is following summer crop production, when conditions become cooler; average temperatures in Minnesota are 30°C high/25°C low in August, but are 14°C high/5°C low in October. Rlv associated with hairy vetch face potentially interacting selection pressures: host plant ecotype and temperature. We do not know the relative contributions of these two factors in determining nodule community assembly, or attendant effects on BNF. In this study, we investigated the relative effects of temperature and hairy vetch ecotype on BNF and nodule Rlv community composition in nodules.

To do so, four ecotypes of hairy vetch were inoculated with soil slurries containing Rlv from one of three locations in Minnesota, then grown at warm or cold temperatures. We pooled all nodules within a temperature x ecotype x soil treatment, used a series of centrifugation steps to separate undifferentiated bacteria from plant material (Burghardt et al., 2018), and used pooled sequencing and bioinformatic analyses to estimate nodule Rlv population differentiation across treatments (Kofler et al., 2011). Allele frequency differences across treatments represent Rlv population differentiation between plants and result from cumulative fitness across three rhizobia life stages: survival before forming nodules, competition between strains for nodulation, and nodule rewards/sanctioning by host plants (Kiers et al., 2003). We sought to answer the following questions: 1) What are the relative contributions of temperature and host plant ecotype, if any, to BNF and Rlv community composition in nodules? 2) Does Rlv community composition in nodules drive effects on plant productivity?

## 2. Methods

### 2.1 Soil collection

To obtain naturalized soil Rlv populations from three distinct locations, we collected soil samples from fields with at least one season of hairy vetch cultivation in the last five years. Samples were collected in August 2022 from three University of Minnesota experimental stations: Roseau (48.88°N, 95.85°W), Rosemount (44.68°N, 93.07°W), and St. Paul (44.99°N, 93.18°W). Ten soil cores were collected to a depth of 15 cm using standard sampling randomization in each field with a 2.5 cm diameter stainless steel soil probe and bucket sterilized with ethanol. Soil cores were combined into one composite sample for each soil origin and stored at 4°C for approximately 3 weeks until experimental use.

### 2.2 Plant genotypes

Hairy vetch ecotypes included the accessions MSP 4045 (V01; Wiering et al., 2018) and Hungvillosa (Brandsaeter et al., 2000), selected for cold tolerance, and the cold-susceptible accessions Purple Bounty (Moore et al., 2020) and AU Merit (Mosjidis, 2002). Seeds were provided by the hairy vetch breeding program at the University of Minnesota in St. Paul, MN.

### 2.3 Seed germination and planting

Seeds of the four ecotypes were surface-sterilized in 90% EtOH for 30 seconds and then in 4% bleach (NaHOCl) solution. Seeds were rinsed repeatedly in sterile water and then germinated in the dark at room temperature for 6 days on sterile petri plates lined with germination paper, moistened with sterile water. Two germinated seeds were aseptically transplanted into a soil microcosm made of two connected Magenta units (Oono et al., 2020) filled with a 1:1 mixture of sand and vermiculite. The soil mix was supplied with N-free nutrient solution from a reservoir made from a third Magenta unit, via cotton wick. Units were thinned to one hairy vetch seedling after one week.

### 2.4 Plant growth

Plants were grown in either a warm (22°C day/18°C night) or cold (15°C day/10°C night) Conviron E8 growth chamber (Conviron, Manitoba, Canada) in St. Paul, MN, with the cold chamber intended to mimic fall temperatures in the upper Midwestern U.S. Experimental chamber conditions were a 12h day/12h night. The experiment was a completely randomized design, with plants re-randomized in the growth chamber weekly. Plants were watered by weight once per week with Hoagland’s N-free nutrient solution, to assess relative growth rates of plants. In Magenta units, which have negligible surface evaporation, water use represents transpiration, which is a useful proxy for leaf area.

### 2.5 Inoculation

For each soil sampling location, 20 g of soil was added to 80 mL of 0.85% NaCl solution to create inoculant slurries. Within each treatment (ecotype x soil origin x temperature), there were four replicate plants, which were inoculated with four replicate soil slurries to ensure that Rlv from across the soil sample were well-represented in inoculants. Slurries were shaken at room temperature at 150 rpm for one hour. After shaking, 1 mL of soil slurry was pipetted into each sterile, sealed, fully self-contained Magenta unit to prevent cross-contamination and colonization from microbes other than those found in soil slurries.

### 2.6 Harvest

Plants were grown for nine weeks to mimic the length of time plants in the field grown in cool, autumn conditions before winter dormancy. At harvest, shoots were separated from roots and dried for 48 hours at 60°C. Shoots were ground to 1 mm and run on a Vario PYRO cube combustion analyzer (Elementar, Germany) to determine percent nitrogen (%N). Nodules were separated from roots, sterilized by soaking for 5 minutes in 4% bleach, and then dessicated using Drierite to obtain nodule dry weight.

### 2.7 DNA extraction

Nodules were pooled for sequencing by treatment (i.e. all replicates of a temperature x ecotype x soil treatment were pooled). Nodules were rehydrated for DNA extraction by soaking in sterile water for one hour and then transferred to 6 mL 0.85% NaCl solution. This mixture was then homogenized using a tabletop tissue homogenizer and centrifuged for 10 min at 400xg to create a pellet enriched for plant tissue and bacteroids and a supernatant enriched for undifferentiated microbial cells (Burghardt et al., 2018). DNA was extracted from supernatants using Qiagen Plant DNeasy Mini Kit following the manufacturers’ instructions.

### 2.8 Nodule Community Analysis

Reads were aligned to the reference genome *R. leguminosarum* bv. *viciae* strain 3841 (Young et al., 2006) using the Call Microbial SNPs application in KBase (Li, 2011). Allele frequencies were calculated across pools using SAMTools (Danecek et al., 2021) and PoPoolation2 (Kofler et al., 2011). To estimate and visualize the contribution of soil origin, ecotype, and temperature to allele frequency shifts across pools, we performed redundancy analysis with the “rda” function in the vegan R package. This allows us to visualize whether soil origin alone determines the mix of Rlv strains in vetch nodules, as well as relative contributions of the plant itself (ecotype) and growing conditions (temperature). We then performed permutations to determine the probability that differences in allele frequencies across pools occurred by chance (“anova” function, 999 permutations). Scripts available at DOI: 10.5281/zenodo.13387709.

### 2.9 Statistical Analyses

Temperature, ecotype, and microbial source effects on plant growth, nodulation, and BNF were measured with Type I ANOVA. Means were separated using Tukey’s HSD tests (p < 0.05). Statistical analysis was performed using R statistical software.

## 3. Results

### 3.1 Temperature and Ecotype Effects on Plant Productivity

As expected, plants accumulated approximately six times more shoot biomass in warm conditions than in cold conditions (*p* < 0.001; **Figure 1**). Ecotype also strongly influenced biomass accumulation (p = 0.001), and there a marginally significant between ecotype and temperature (*p* = 0.055). Under cold conditions, the ecotype AU Merit accumulated more biomass than any other ecotype. Under warm conditions, AU Merit accumulated more biomass than Purple Bounty, but a statistically equivalent amount to Hungvillosa and MSP 4045. While AU Merit is considered “cold-susceptible” due to low freezing tolerance, this study did not test freezing tolerance, which is the primary breeding target for cold-tolerant ecotypes. AU Merit was bred for high biomass accumulation (Mosjidis, 2002), which this study suggests it can still achieve under relatively low temperature conditions (15°C day/10°C night).

**Figure 1.**
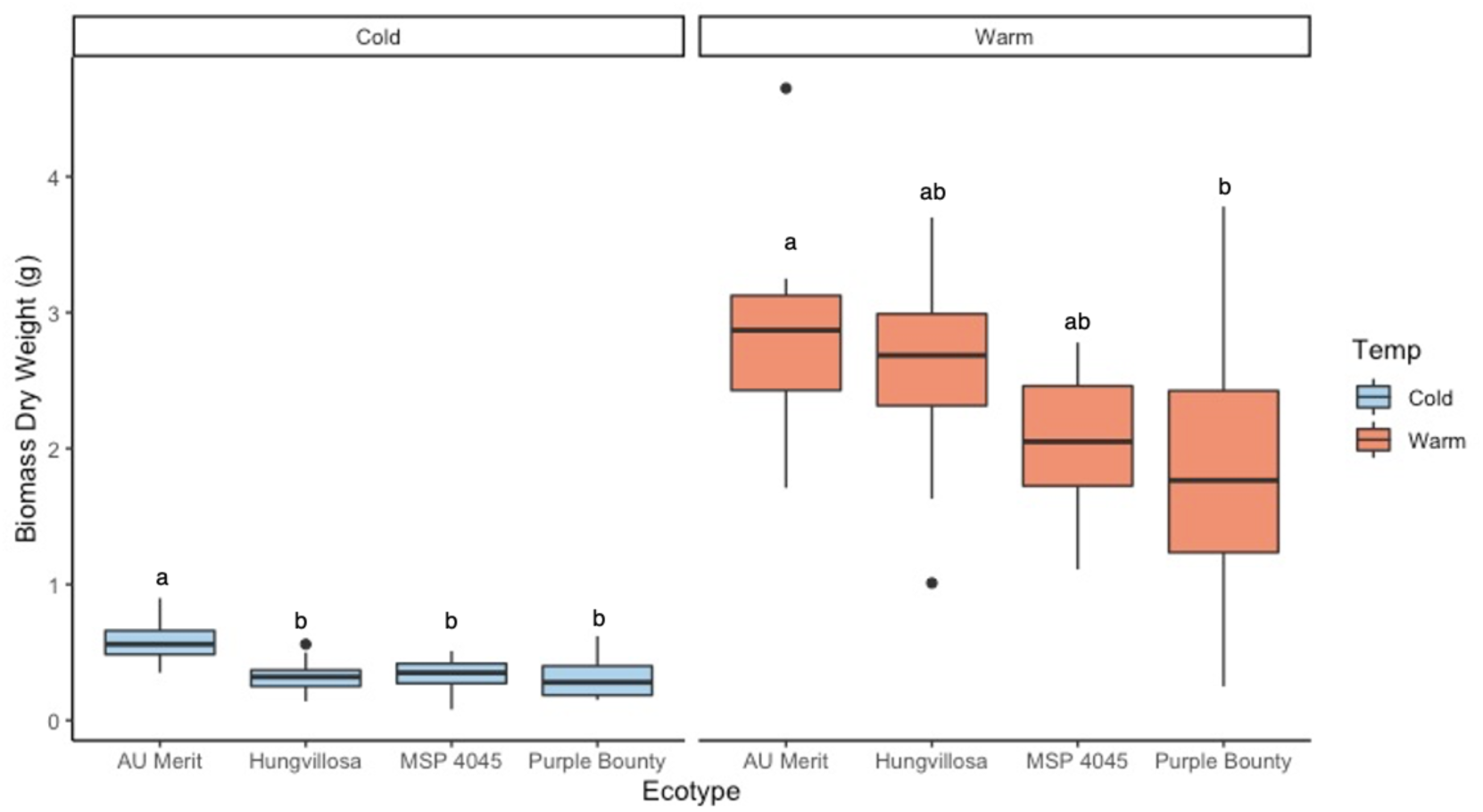
Shoot biomass dry weight per plant of four hairy vetch ecotypes grown in warm or cold conditions. Letters indicate significant difference between means within a temperature treatment. Means were separated by Tukey’s HSD (*p* < 0.05).

### 3.2 Temperature and Ecotype Effects on Nodulation

Temperature and ecotype interacted to affect nodule dry mass per plant (*p* = 0.02). Cold conditions reduced nodule dry weight per plant compared to warm conditions (*p* < 0.001; **Figure 2**), and ecotype also drove nodule dry mass per plant (*p* < 0.001). Mirroring shoot biomass results, AU Merit plants had greater nodule mass per plant than other ecotypes at low temperatures. At warm temperatures, AU Merit again had the greatest nodule mass, and Purple Bounty had the least. However, at both temperatures, Purple Bounty had a greater return on nodule construction cost (shoot mass/nodule mass) than AU Merit (**Supplemental Figure 1).**

**Figure 2.**
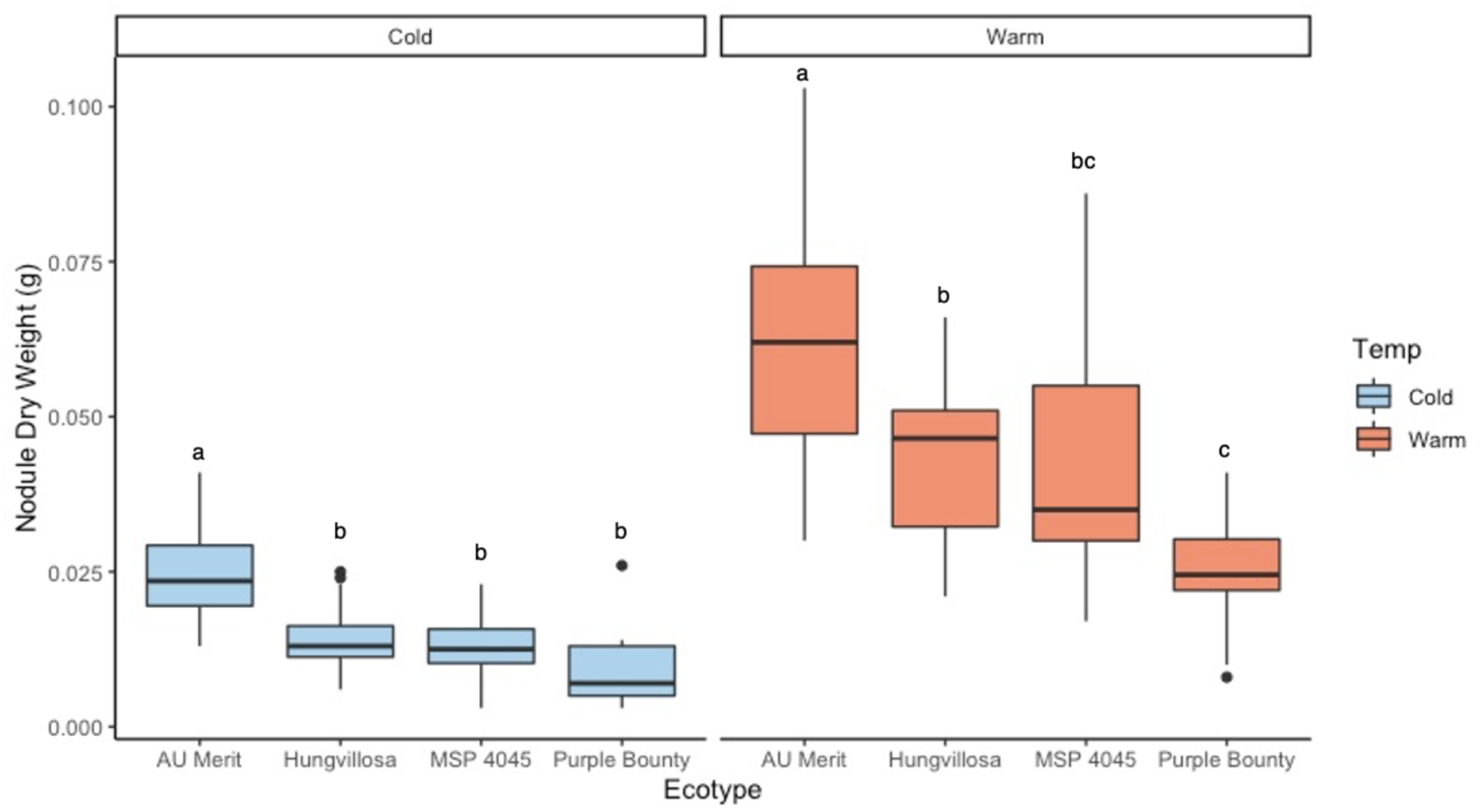
Nodule dry weight per plant of four hairy vetch ecotypes grown in warm or cold conditions. Letters indicate significant difference between means within a temperature treatment. Means were separated by Tukey’s HSD (*p* < 0.05).

### 3.3 Temperature and Ecotype Effects on Biological Nitrogen Fixation

Total shoot N was lower in cold conditions than warm conditions, due to the combination of less BNF and less shoot biomass accumulation in the cold (*p* < 0.001; **Figure 3**). Total shoot N is calculated as the %N present in shoot biomass, multiplied by total shoot biomass, which can be used to estimate N contributions from hairy vetch plants when incorporated into the soil. Ecotype also strongly influenced total N per plant (*p* = 0.001), and there was only a weak interaction between ecotype and temperature (*p* = 0.03). Soil origin had no effect on total N per plant (*p* > 0.05), indicating that differences among rhizobia populations overall did not drive differences in BNF. Nodule mass was a stronger predictor of total plant N under cold conditions than warm conditions (R^2^ = 0.84 and 0.59 respectively, *p* = 0.01, **Figure 4**).

**Figure 3.**
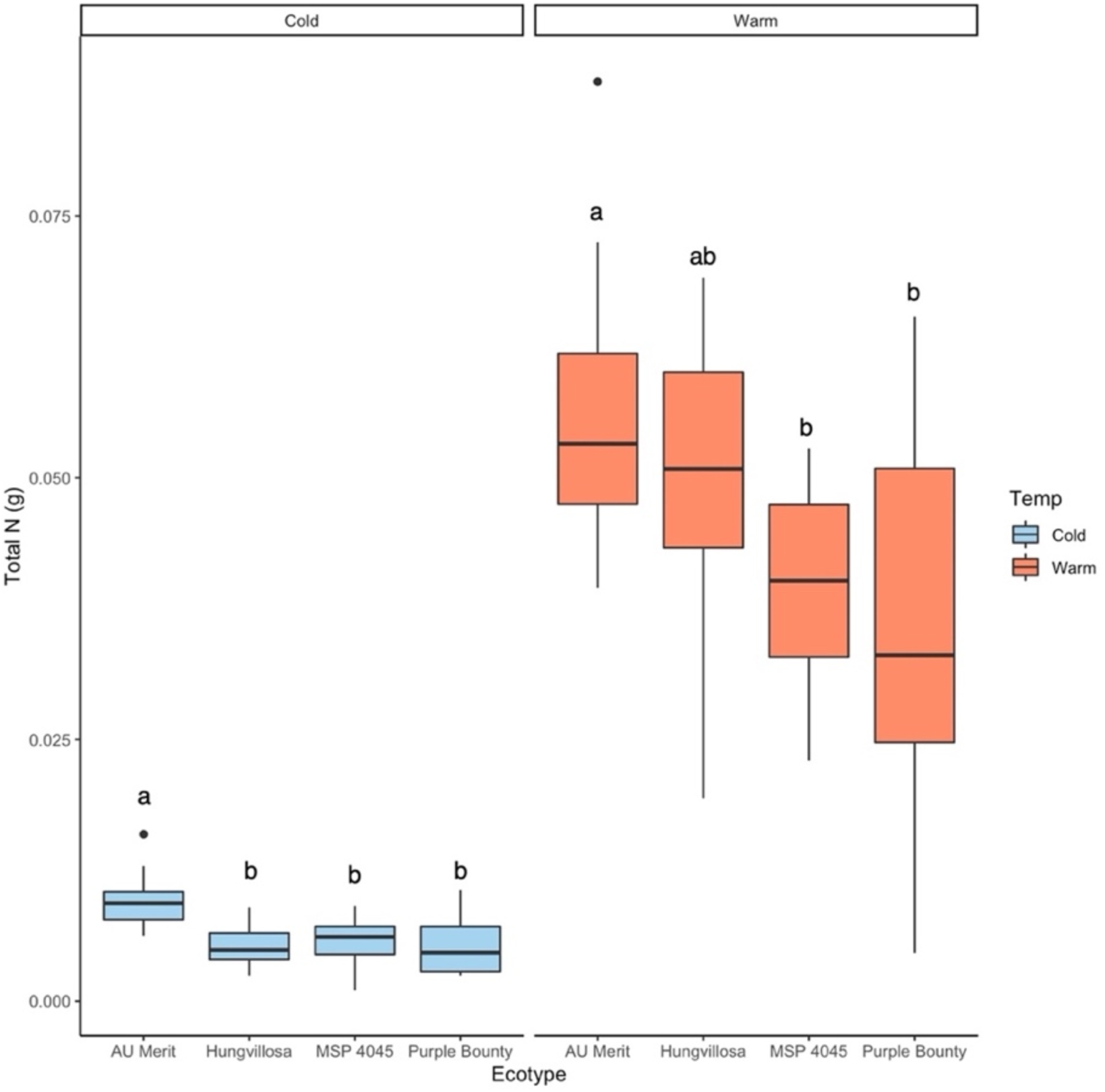
Total shoot N of four hairy vetch ecotypes grown in warm or cold conditions. Letters indicate significant difference between means within a temperature treatment. Means were separated by Tukey’s HSD (*p* < 0.05).

**Figure 4.**
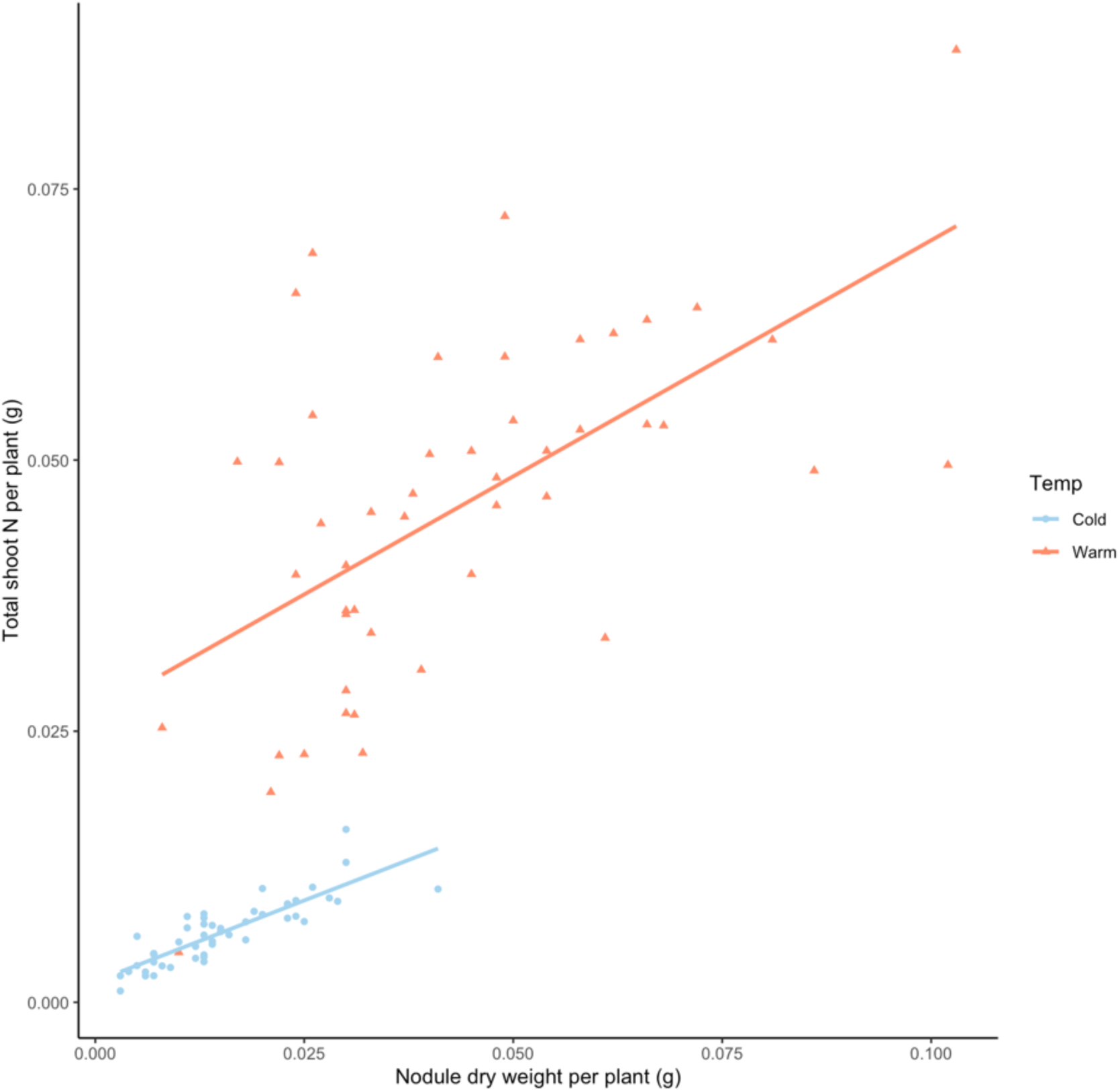
Nodule dry weight regressed against total plant N for four vetch ecotypes at two temperatures. Each point represents one hairy vetch plant. Nodule mass is a stronger predictor of total plant N under cold conditions than warm conditions (R^2^ = 0.84 and 0.59 respectively, *p* = 0.01).

Under cold conditions, the ecotype AU Merit accumulated more shoot N than any other ecotype. Under warm conditions, AU Merit accumulated more shoot N than Purple Bounty and MSP 4045, but a statistically equivalent amount to Hungvillosa. While AU Merit is considered “cold-susceptible,” meaning that it has relatively low winter survival due to low freezing tolerance (Wiering et al., 2018), it still achieved relatively high N accumulation under cool conditions (15°C day/10°C night).

### 3.4 Temperature and Ecotype Effects on Nodule Community Composition

One potential explanation for the phenotypic patterns we observed could be that plants were interacting with different communities of rhizobia. Soil origin explained 18.8% of the variance in overall nodule community composition (*p* = 0.02; **Figure 5)** suggesting that soil origins diverge Rlv populations. Given our experimental design, we had limited power to detect ecotype effects. However, across all soil origins, ecotype explained 10.3% of variance (*p* = 0.59) and temperature explained 3.5% of variance (*p* = 0.46). To assess if temperature altered rhiozbial composition within soil origin treatments, we analyzed each soil origin separately. Intriguingly, we found that the importance of temperature depended on the microbial source (**Supplemental Figure 2**). At Roseau, temperature explained 17.1% of variance (*p* = 0.1) and at Rosemount, temperature explained 31.5% of variance (*p* = 0.03). However, at St. Paul, Temperature explained only 7.5% of variance (*p* = 0.69). This suggests that the effect of temperature varies among soil rhizobia populations, possibly due to effects not explored in this study such as planting history or soil pH.

**Figure 5.**
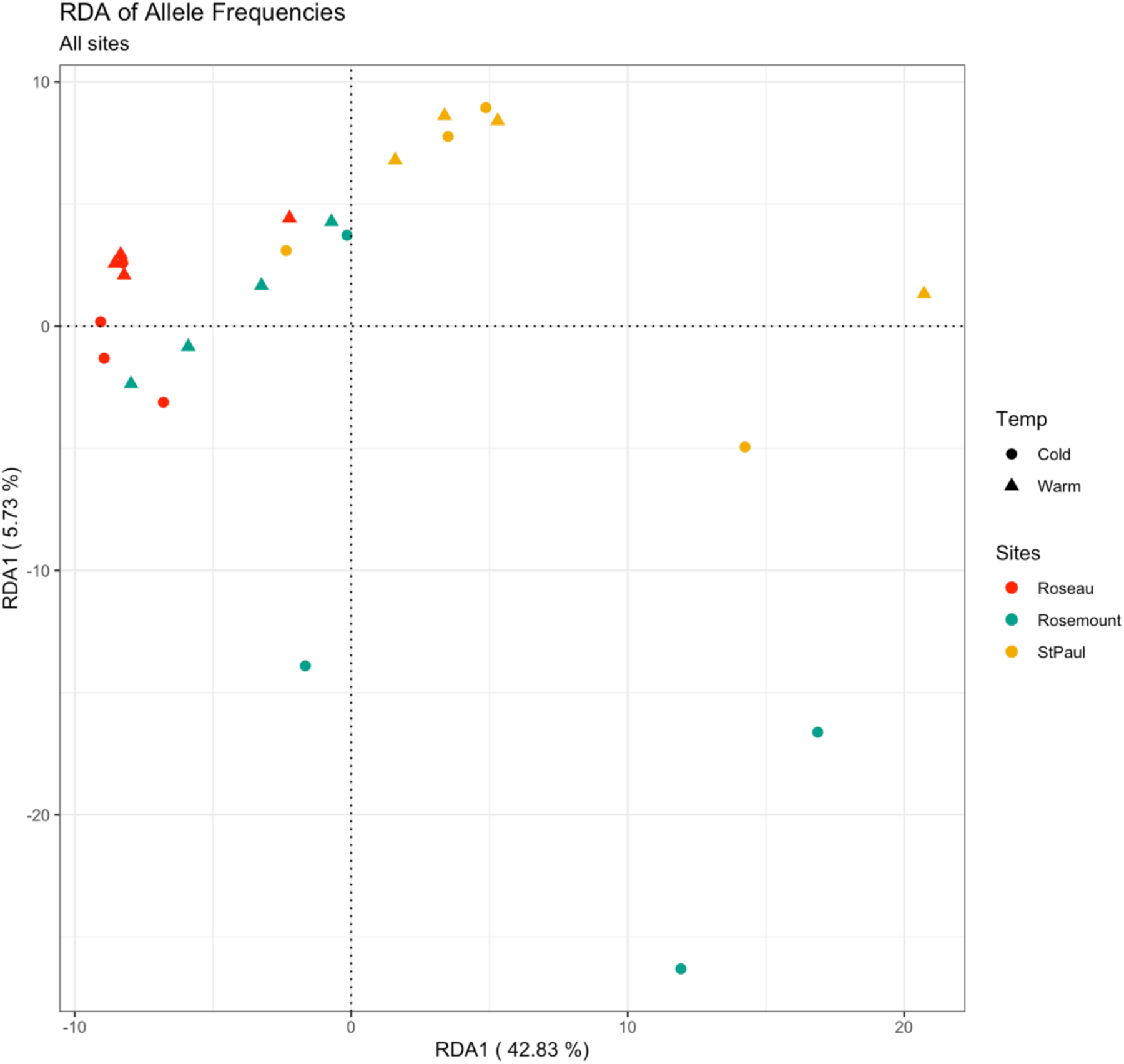
RDA analysis shows that soil origin affected nodule community characteristics, explaining 19.3% of variance (*p* = 0.01). Host ecotype explained 9.8% of variance (*p* = 0.64), and temperature explained 3.3% of variance (*p* = 0.51).

## 4. Discussion

### 4.1 Temperature and ecotype drive hairy vetch productivity and BNF

We found that low temperatures reduced hairy vetch productivity and BNF across multiple ecotypes, including two ecotypes bred for winter hardiness. Our results are supported by field and growth chamber experiments showing decreased biomass accumulation and BNF in hairy vetch at temperatures below 20°C (Thurston et al., 2022). In this study, plants grew for nine weeks under cold conditions, reflecting the period when vetch must accumulate biomass before winter dormancy. In northern zones such as the Upper Midwestern U.S., mean air temperatures of 10° – 15°C begin as early as October, when hairy vetch is frequently sown following summer cash crop production. Our results reflect the significant challenge of achieving meaningful biomass and N accumulation under cold fall conditions before winter dormancy.

Our results also showed variation in shoot biomass accumulation among ecotypes at warm temperatures, which is contrary to other hairy vetch growth chamber studies (N. V. Mothapo et al., 2013). This is likely due to our study’s use of ecotypes bred for specific plant traits. For example, the ecotype AU Merit accumulated more biomass than other ecotypes under both temperature conditions. The ecotype was originally bred for high biomass accumulation but not freezing tolerance (Mosjidis, 2002), which supports these results. AU Merit’s relatively strong performance under cold conditions compared to winter-hardy ecotypes reflects evidence that ecotypes adapted to colder regions, such as Arctic sedge species, may have reduced plasticity in response to changing environmental conditions because those ecotypes evolved under more constrained environmental conditions (Curasi et al., 2019).

Producers who use hairy vetch as a cover crop aim to introduce new fixed nitrogen to the system. Currently, the ecotype that this study showed to provide the most nitrogen under cold conditions, AU Merit, is unlikely to survive freezing winter conditions. This reflects a challenge in breeding winter annual crops, wherein there may be a tradeoff between high biomass accumulation and freezing tolerance. Yet, both traits are critical for a cover crop to deliver sufficient nitrogen to a field. In fact, this challenge is so acute that in Minnesota, high fall vigor of hairy vetch is a negative predictor of spring vigor (Kucek et al., 2019), and spring vigor ultimately determines the degree of biomass incorporated into the soil. Both the most and least vigorous plants in the fall tended to underperform in the spring, possibly because they either had not developed enough resources to survive the winter or because they grew large vegetative architecture (at the expense of root reserves) that was vulnerable to winter damage. However, the smallest plants in the fall also underperformed in the spring, which suggests there is an ideal biomass quantity for vetch to accumulate in the fall that will allow for winter survival and high biomass accumulation in spring. Future breeding efforts must balance these tradeoffs. The results of this study suggest that higher biomass ecotypes, such as AU Merit, may contribute valuable genetic stock to breeding programs, to continue to improve biomass accumulation in freezing-tolerant varieties which fix less nitrogen such as MSP 4045.

### 4.2 Temperature and ecotype effects on nodulation

Hairy vetch nodule mass per plant followed a similar pattern as biomass accumulation, wherein cold temperatures reduced nodule mass, and ecotype also affected nodule mass similarly to shoot biomass. The treatment with the highest nodule mass under cold conditions (AU Merit) had a similar nodule mass to the nodules of at least one ecotype treatment under warm conditions (Purple Bounty). Mean nodule size can vary among host plants from different populations within a species (Wendlandt et al., 2019). A plant can sanction poor nitrogen-fixers by cutting off carbon resources to individual nodules that fix less than others on the same plant (Kiers et al., 2003). The study by Wendlandt et al. suggests that nodules size can still vary across populations of host plants even when those plants show evidence of similarly robust sanctions. Future experiments could explore whether greater nodule mass in AU Merit represents stricter sanctions by the host plant or an intrinsic property of the ecotype to invest in larger nodules regardless of BNF output.

### 4.3 Soil, ecotype, and temperature effects on nodule community composition

It is well-documented that soil origin drives rhizosphere microbiome assembly across many plant species (Brown et al., 2020; Edwards et al., 2015; Liu et al., 2019; Wagner et al., 2016). Soil origin has also been shown to drive nodule community composition in hairy vetch, both in experiments where a soil slurry was used as a rhizobial inoculant (N.V. Mothapo et al., 2013) and when plants trapped rhizobia directly form soil (Zhang et al., 2020), which aligns with the results of the study. Soil origin did not affect hairy vetch biomass, shoot nitrogen, or nodule mass, but did affect nodule community composition. This result underscores the existence of highly variable endogenous rhizobia populations within and across fields, and in this study, soil effects on rhizobia populations outpaced either ecotype or temperature effects. In previous studies, plant genotype has been shown to structure nodule community (Bourion et al., 2018; Burghardt et al., 2018). In contrast, hairy vetch may exert less consistent selective pressure on rhizobia due to its high genetic variability within an ecotype, relative to other studied legumes. Some evidence suggests that domesticated species may be inferior in their ability to regulate microbiomes when compared to wild relatives (Bouffaud et al., 2012; Bulgarelli et al., 2015; Pérez-Jaramillo et al., 2016), but few studies have investigated a semi-domesticated crop like hairy vetch.

### 4.4 Nodule community assembly does not drive plant productivity

Rhizobia community composition varied only with soil origin, which appeared to have little effect on shoot biomass or nitrogen accumulation. Agricultural fields can host a broad range of rhizobia, which vary in traits such as nitrogen fixation rate and, consequently, benefit to the plant. There is evidence, based on all legumes studied so far, that legumes can selectively reward rhizobia that fix nitrogen more efficiently (Kiers et al., 2003; Oono et al., 2011; Regus et al., 2017), and that particular plant genotypes within a species can select more beneficial rhizobia than other genotypes within the same species (Burghardt et al., 2018). Thus, we originally hypothesized that plants with greater shoot nitrogen would associate with superior rhizobia strains than plants with less shoot nitrogen. However, nodule population composition did not correlate with plant productivity. Unless we develop methods whereby inoculant strains can dominate nodules in the field, the most productive path forward to improve nitrogen accumulation in vetch may be to focus on plant breeding, rather than harnessing specific associations with rhizobia.

## 5. Conclusions

In summary, we examined the effects of temperature and host plant ecotype on hairy vetch BNF and Rlv community composition in nodules. Our main findings were that (1) hairy vetch ecotype drove BNF and nodule formation at warm and cold temperatures, (2) soil origin drove nodule community composition, rather than temperature or hairy vetch ecotype, and (3) nodule community composition did not correlate with plant productivity. These findings expand our understanding of temperature’s role in BNF, nodule formation, and nodule community composition. Taken together, our results suggest that BNF could be improved in hairy vetch at low temperatures through breeding for improved biomass accumulation and nitrogen fixation in host plants, rather than by focusing on host plant selection of beneficial rhizobia.

## CRediT authorship contribution statement

**Rebecca Fudge:** Conceptualization, Methodology, Formal Analysis, Investigation, Data Curation, Writing – Original Draft, Writing – Review & Editing, Visualization, Funding Acquisition. **Paula Gardner:** Methodology, Software, Validation, Formal Analysis, Data Curation, Writing – Review & Editing. **R. Ford Denison:** Conceptualization, Writing – Review & Editing, Supervision. **Liana Burghardt:** Conceptualization, Writing – Review & Editing, Supervision. **Julie Grossman:** Conceptualization, Resources, Writing – Review & Editing, Supervision, Project Administration, Funding Acquisition.

## Acknowledgments

This study was funded by U.S. Department of Agriculture (USDA), National Institute of Food and Agriculture (NFIA) Organic Transitions Program (1017066), North Central Sustainable Agriculture Research and Education Graduate Student Grant (GNC20-300), Minnesota Department of Agriculture Specialty Crop Block Grant (2021-HR133-11), and USDA NIFA Predoctoral Fellowship (1030772). These findings should not be construed to represent any agency determination or policy.

