## Supplemental Figure 1 for "Temperature and host plant ecotype drive nitrogen fixation, but not nodule community composition, in hairy vetch"

Supplemental Figures

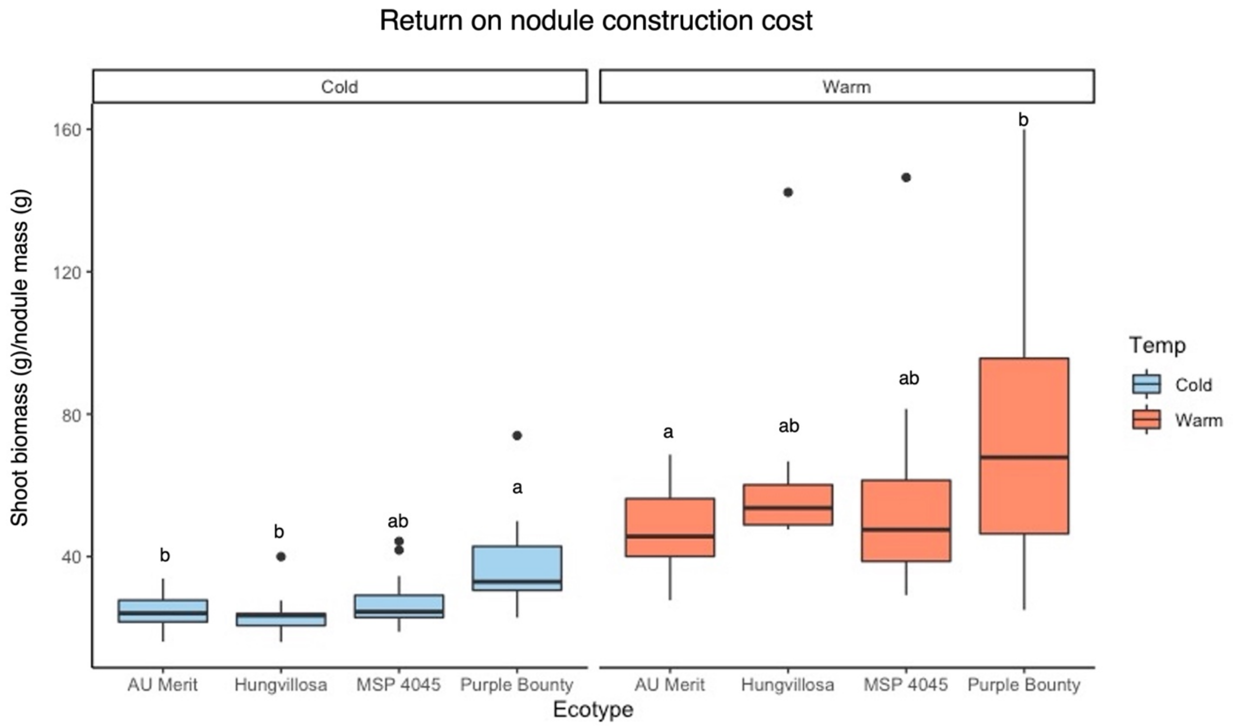


**Supplemental Figure 1** Return on nodule construction cost is calculated as shoot biomass/nodule mass per plant. Purple Bounty had a higher return on nodule construction cost (shoot biomass/nodule mass) than AU Merit at both warm and cold temperatures. Letters indicate significant difference between means within a temperature treatment. Means were separated by Tukey's HSD ($p < 0.05$).

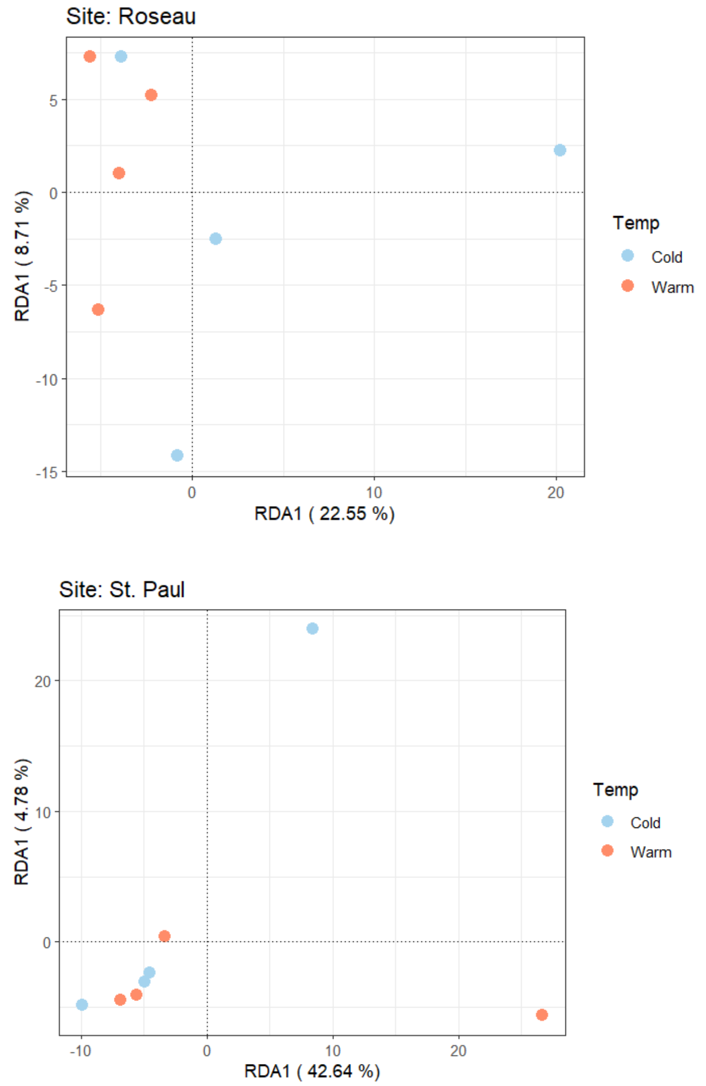


**Supplemental Figure 2** RDA of nodule allele frequencies, partitioned by soil origin. The effect of temperature on nodule community composition varies across soil origins. Roseau variance explained = 17.1% ($p = 0.1$), Rosemount variance explained 34.3% ($p = 0.03$), St. Paul variance explained = 7.5% ($p = 0.69$).
